# AKAP79/150 coordinates leptin-induced PKA activation to regulate K_ATP_ channel trafficking in pancreatic β-cells

**DOI:** 10.1101/2020.11.25.397059

**Authors:** Veronica A. Cochrane, Zhongying Yang, Mark Dell’Acqua, Show-Ling Shyng

## Abstract

The adipocyte hormone leptin regulates glucose homeostasis both centrally and peripherally. A key peripheral target is the pancreatic β-cell, which secretes insulin upon glucose stimulation. Leptin suppresses glucose-stimulated insulin secretion by promoting trafficking of K_ATP_ channels to the β-cell surface, which increases K^+^ conductance and causes β-cell hyperpolarization. Here we investigate the signaling mechanism underlying leptin-induced K_ATP_ channel translocation with a focus on protein kinase A (PKA). Using FRET-based PKA activity reporters, we show that leptin increases PKA activity at the cell membrane via a signaling pathway involving NMDA receptors, CaMKKβ and AMPK. Genetic knockdown and rescue experiments reveal that leptin activation of PKA requires tethering of PKA to the membrane-targeted PKA-anchoring protein AKAP79/150. Interestingly, disrupting protein phosphatase 2B (PP2B) anchoring to AKAP79/150, known to elevate basal PKA signaling, increases surface K_ATP_ channels. Our findings uncover a novel role of AKAP79/150 in coordinating leptin and PKA signaling to regulate β-cell function.

## Introduction

Pancreatic β-cells produce and secrete insulin making them critical for maintaining glucose homeostasis. In order to secrete insulin in a timely and controlled manner, β-cells must interpret and respond to a myriad of physiological stimuli and hormones. One such hormone is the adipocyte-derived hormone leptin, which regulates serum insulin levels by suppressing glucose-stimulated insulin secretion (GSIS) from β-cells^1–5^. Leptin does so by promoting trafficking of ATP-sensitive potassium (K_ATP_) channels to the β-cell membrane^6–10^. This increased surface expression of K_ATP_ channels increases the total K_ATP_ channel conductance thereby causing membrane hyperpolarization and reducing β-cell electrical activity. Our studies to date have shown that leptin-induced K_ATP_ channel trafficking involves the following series of signaling events^7–10^. Leptin stimulates Src kinase to phosphorylate and potentiate NMDA receptor (NMDAR) activity, resulting in enhanced Ca^2+^ influx that activates calcium/calmodulin-dependent kinase kinase β (CaMKKβ), which then phosphorylates and activates AMP-activated protein kinase (AMPK). Downstream of AMPK actin depolymerization occurs and results in increased trafficking of K_ATP_ channels to the plasma membrane^7–10^. We have shown previously that actin remodeling and subsequent K_ATP_ channel trafficking following AMPK activation requires protein kinase A (PKA). However, it remains elusive whether PKA functions as an active or permissive player in leptin signaling.

PKA is a serine/threonine kinase consisting of two regulatory subunits and two catalytic subunits. Binding of cAMP to the regulatory subunits causes a conformational change that releases the autoinhibition of the catalytic subunits to activate PKA^11,12^. Due to the ubiquity of PKA in mammalian cells and the promiscuity of its catalytic subunits, PKA is involved in a multitude of cellular processes. In pancreatic β-cells PKA has been identified by several studies as a positive regulator of insulin granule trafficking and exocytosis^13–17^. However, evidence that PKA promotes K_ATP_ channel trafficking suggests that PKA is also involved in signaling events that lead to inhibition of insulin secretion^7,18^. These findings illuminate the complexity of β-cell signaling and raise the question of how cellular signaling networks are coordinated to fine-tune insulin secretion. It is well established that PKA interacts with a family of scaffolding proteins termed A-kinase anchoring proteins (AKAPs). Different AKAPs direct PKA activity towards specific cell signaling machinery by anchoring PKA and other signaling molecules to distinct subcellular locations^19^. Several AKAPs have been identified as having roles in regulating insulin secretion, but the mechanisms by which they do so remain poorly understood^20–25^.

In this study we show that leptin increases PKA activity at the β-cell membrane, but not in the cytoplasm, through our previously identified pathway involving NMDAR, CaMKKβ, and AMPK. Moreover, we find that anchoring of PKA by AKAP79/150 is necessary for leptin-mediated K_ATP_ channel trafficking and that overexpression of an AKAP79/150 mutant associated with increased basal PKA activity due to a loss of protein phosphatase 2B (PP2B) binding recapitulates the effects of leptin. Additionally, we present evidence that leptin locally increases concentrations of the PKA activator cAMP near AKAP79/150 expressed at the cell membrane but not in proximity to AKAP18δ, which is primarily found in the cytoplasm. These findings reveal a novel function of the PKA-AKAP79/150 signaling complex for orchestrating leptin signaling to regulate KATP channel surface expression and thus β-cell excitability.

## Results

### Leptin increases PKA activity near the plasma membrane

We have previously shown that leptin promotes trafficking of KATP channels to the plasma membrane, causing increased K^+^ conductance and cell hyperpolarization in rat insulinoma INS-1 832/13 cells as well as primary mouse and human β-cells^7–10^. Our prior work revealed that a critical event for this process is actin remodeling^7^, which presumably allows vesicles containing potassium channels to translocate and insert into the β-cell membrane^26–29^. Furthermore, we found that the ability of leptin to remodel the actin cytoskeleton and increase surface KATP channel density could be blocked by the protein kinase A (PKA)-specific inhibitor peptide (PKI) suggesting that PKA is essential to this signaling mechanism^7^. However, whether leptin stimulation leads to direct activation of PKA remains unknown. To determine whether leptin signaling activates PKA we monitored PKA activity using the FRET-based PKA activity reporter A-kinase activity reporter 4 (AKAR4)^30^. AKAR4 contains a PKA substrate motif that is phosphorylated by PKA causing the sensor to undergo a conformational change to increase FRET detected as an increased YFP acceptor/CFP donor emission ratio in response to excitation of the CFP donor. An AKAR4 targeted to the plasma membrane with a farneyslation motif (CAAX; Fig. 1A) as well as an AKAR4 targeted to the cytoplasm by a nuclear export signal (NES; Fig. 1B) were both tested. INS-1 832/13 cells expressing either AKAR4-CAAX or AKAR4-NES were treated with leptin (100 nM) or vehicle. In order to control for the expression level of AKAR4 between cells the potent PKA activator forskolin was administered at the end of each experiment and cells that responded robustly to forskolin with a 10-15% increase in FRET were chosen for analysis. For quantification purposes, FRET traces were normalized to the maximal forskolin response (Fig. 1C,D) and then analyzed for area under the curve (AUC) during the treatment period with vehicle or leptin. In AKAR4-CAAX expressing cells, leptin treatment led to a 6.37-fold increase in PKA activity (p<0.0005, n=24) compared to vehicle treated cells (n=13) (Fig. 1E). In contrast, AKAR4-NES expressing cells showed no change in PKA activity between vehicle (n=9) and leptin (n=10) treatments (Fig. 1F). To confirm that the increase in PKA activity in AKAR4-CAAX expressing cells was due to leptin and not an experimental artifact we tested the effect of various leptin concentrations compared to inactive boiled leptin. While we found that leptin at concentrations of 10 nM and 100 nM increased PKA activity, boiled 100 nM leptin did not (Fig. 1G). This result indicates that leptin does indeed increase PKA activity and that this increase is spatially restricted to the plasma membrane.

**Figure 1.**
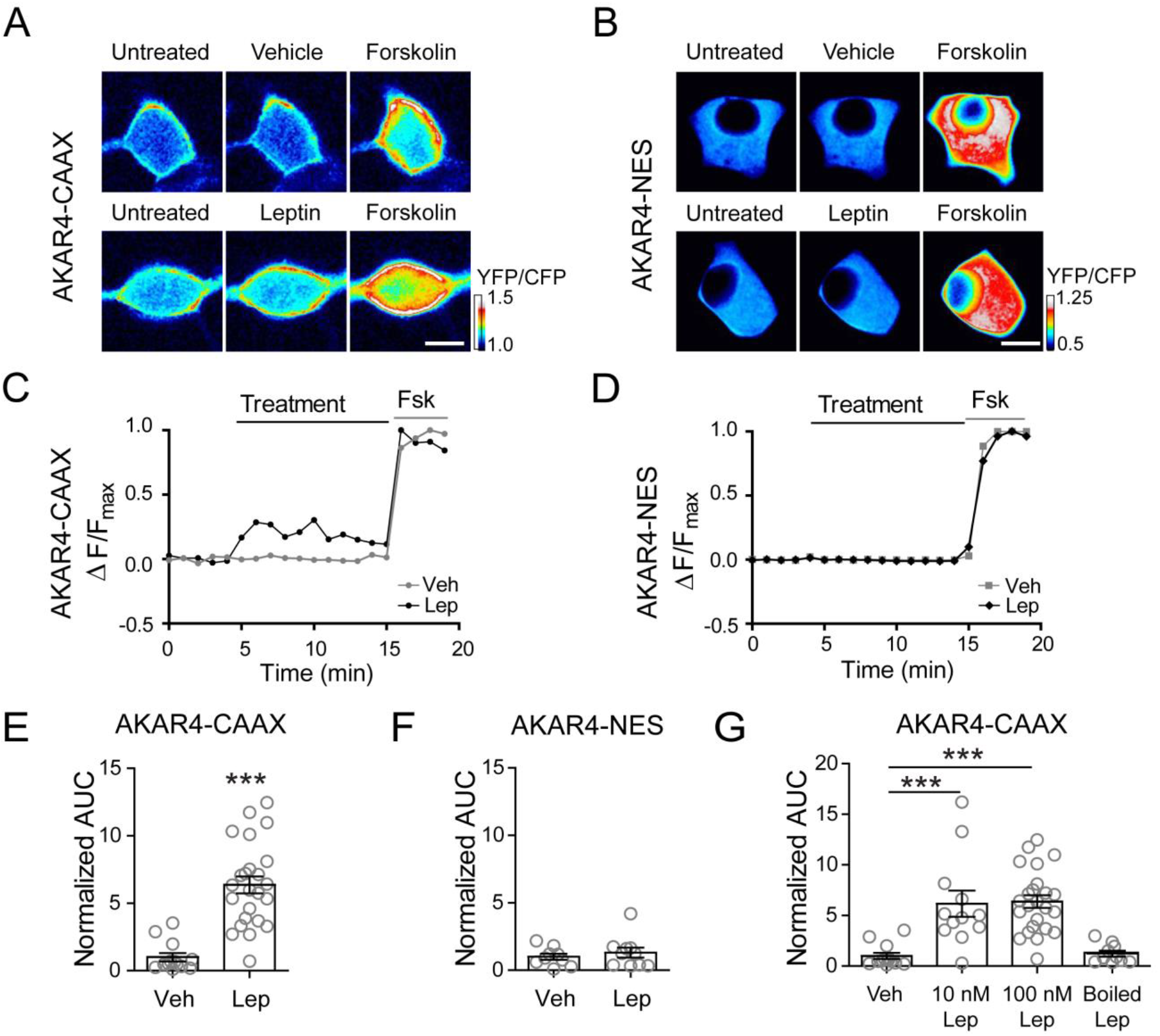
Leptin increases PKA activity. A, B) Ratiometric images of INS-1 832/13 cells transfected with the FRET-based PKA activity reporter A-kinase activity reporter 4 (AKAR4), which has been targeted to the plasma membrane with a farnesylation motif (AKAR4-CAAX) or to the cytoplasm with a nuclear export signal (AKAR4-NES). Cells were treated with vehicle or leptin (100 nM) followed by the robust PKA activator forskolin (20 μM). *Scale bar, 5 μm*. C, D) FRET traces of the cells in (A, B) normalized to the maximal forskolin response. E) Group analysis of AKAR4-CAAX cell traces. Graph shows the fold-change in area under the curve (AUC) normalized to vehicle treatment (n=13). 100 nM leptin (n=24) was used for these experiments. ***p<0.0001 by unpaired student’s t-test. F) Group data of AKAR4-NES expressing cells treated with vehicle (n=9) or 100 nM leptin (n=10). G) Group analysis for AKAR4-CAAX cells treated with vehicle (n=13), 10 nM leptin (n=12), 100 nM leptin (n=24), or boiled 100 nM leptin (n=10). ***p<0.001 by one-way ANOVA followed by a post-hoc Dunnett’s multiple comparison test. In all figures, circles represent individual cells except as otherwise specified.

### Leptin signals via the NMDAR-CaMKKβ-AMPK axis to activate PKA

Next, we sought to determine the signaling mechanism that underlies leptin activation of PKA. Our previous studies have shown that the leptin signaling pathway leading to increased K_ATP_ channel trafficking involves potentiation of NMDAR activity and Ca^2+^ influx to activate CaMKKβ; this results in phosphorylation and activation of AMPK, which is followed by PKA-dependent actin depolymerization^7–10^. Thus, a logical hypothesis is that leptin activates PKA via the NMDAR-CaMKKβ-AMPK signaling axis. To test this, we implemented the membrane targeted PKA activity sensor AKAR4-CAAX in conjunction with pharmacological reagents. As shown in Fig.2A, inhibiting the initial potentiation of NMDARs with the competitive NMDAR antagonist D-APV (50 μM) or preventing the ensuing Ca^2+^ influx with the Ca^2+^ chelator BAPTA (10 μM) reduced leptin activation of PKA by 2.5-fold (n=12) and 5-fold (n=11) respectively, levels that were not significantly different from the vehicle control. Consistent with these findings, inhibiting CaMKKβ with STO-609 (1 μM) also prevented leptin from increasing PKA activity (n=9). Finally, the role of AMPK for leptin activation of PKA was tested using Compound C (CC, also known as Dorsomorphin, 1 μM). Blocking AMPK activity with CC (n=15) greatly diminished the effect of leptin such that PKA activity levels resembled those of vehicle treated cells. These results support the notion that PKA is activated downstream of AMPK in the leptin signaling cascade.

**Figure 2.**
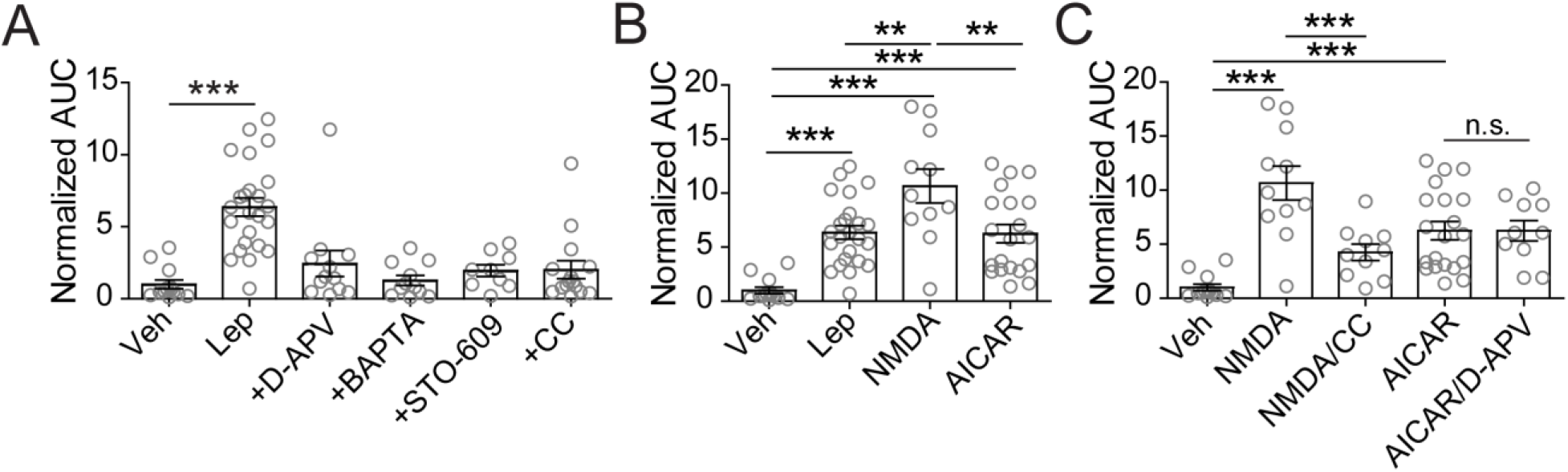
Leptin activates PKA via the NMDAR-CaMKKβ-AMPK signaling cascade. INS-1 832/13 cells were transfected with AKAR4-CAAX followed by various treatments. A) Activation of PKA in response to 100 nM leptin (n=24) alone or in the presence of the NMDAR inhibitor D-APV (50 μM; n=12), the Ca^2+^ chelator BAPTA (10 μM; n=11), the CaMKKβ inhibitor STO-609 (1 μM; n=9), or the AMPK inhibitor Compound C (CC, 1 μM; n=15). Treatments were compared to vehicle control (n=13). ***p<0.001 as determined by one-way ANOVA followed by a post-hoc Dunnett’s multiple comparison test. B) Effects of NMDAR co-agonists NMDA/glycine (100 μM/100 μM; n=11) and the AMPK activator AICAR (500 μM; n=20) on PKA activity. C) PKA activity in response to NMDAR activation by NMDA/glycine in the absence or presence of the AMPK inhibitor CC (1 μM; n=10), and to AMPK activation by AICAR in the absence or presence of the NMDAR inhibitor D-APV (50 μM; n=10). **p<0.01, ***p<0.001 by one-way ANOVA followed by a post-hoc Tukey’s multiple comparison test unless stated otherwise.

To further corroborate the above findings, we tested whether pharmacological activation of NMDAR or AMPK could increase PKA activity. Treating AKAR4-CAAX expressing cells with the NMDAR co-agonists NMDA (100 μM) and glycine (100 μM) or the AMPK agonist AICAR (500 μM) significantly increased PKA activity (Fig. 2B). While direct activation of AMPK caused a 6.25-fold increase (p<0.001, n=20) over vehicle that was equivalent to leptin treatment, NMDAR co-agonists caused a 10.66-fold increase (p<0.001, n=11) compared to vehicle, which was 1.67 times (p<0.01, n=11) greater than that of leptin. This suggests that the direct activation of NMDARs may have caused a greater extent of NMDAR activity and consequently more PKA activity. To confirm that NMDARs signal through the leptin pathway involving AMPK we treated cells with the NMDAR co-agonists NMDA and glycine in the presence of the AMPK inhibitor CC (1 μM) and found that CC significantly reduced the effect of NMDAR activation on PKA (n=10, p<0.001) (Fig. 2C). On the contrary, the effects of the AMPK agonist AICAR were not blocked by the NMDAR inhibitor D-APV (50 μM) indicating that NMDAR lies upstream of AMPK. Collectively, these findings demonstrate that leptin signaling through the NMDAR-CaMKKβ-AMPK cascade increases PKA activity.

### Leptin signaling via PKA requires AKAPs

During the live cell PKA activity imaging experiments it was apparent that leptin increased PKA activity at the cell membrane but not throughout the cytoplasm. This suggests that leptin activates a subset of cellular PKA that is localized near the plasma membrane. It is widely documented that a high level of regulation and specificity of PKA signaling is maintained by a family of scaffolding proteins known as A-kinase anchoring proteins (AKAPs), which target PKA and its signaling partners to distinct subcellular regions thereby creating PKA signaling microdomains and nanodomains^19,31,32^. To test if AKAPs are involved in targeting PKA to the cell membrane for leptin signaling we used the PKA-AKAP interaction disruptor peptide st-Ht31 (50 μM)^33^. This peptide binds the regulatory subunits of PKA and prevents PKA from binding AKAPs. We first introduced st-Ht31 to INS-1 832/13 cells expressing AKAR4-CAAX and monitored PKA activity in response to various stimuli (Fig. 3A). Disrupting the PKA-AKAP interaction with st-Ht31 led to a significant decrease in PKA activation by leptin (3.6-fold decrease, p<0.05, n=7), NMDAR (6.4-fold decrease, p<0.01, n=8), and AMPK (3.4-fold decrease, p<0.001, n=10). These experiments showed that activation of PKA at the cell membrane by leptin signaling requires AKAPs.

**Figure 3.**
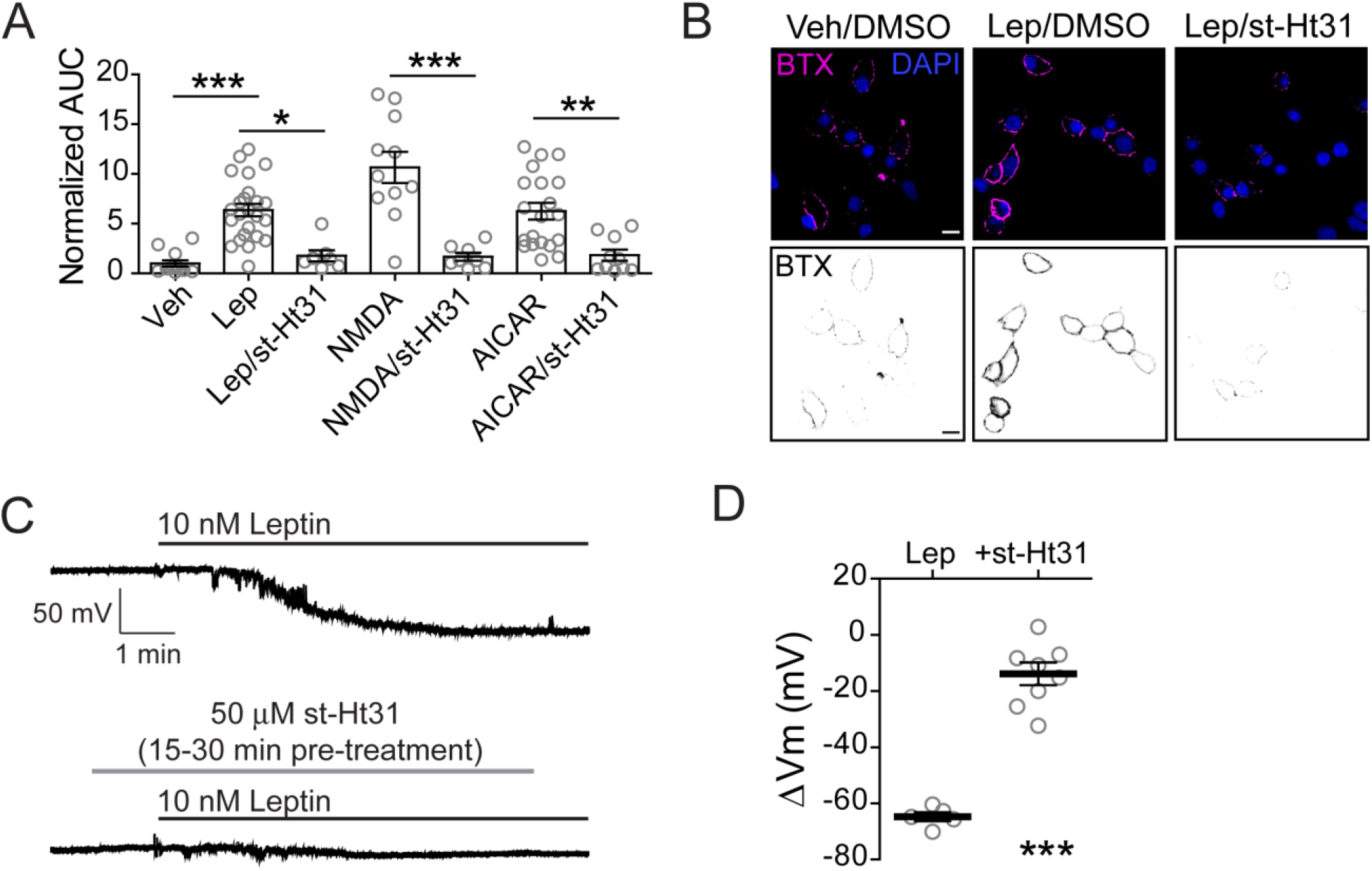
Leptin signaling through PKA requires an A-kinase anchoring protein (AKAP). A) Effects of the PKA-AKAP interaction disruptor peptide st-Ht31 (50 μM) on PKA activity. Group FRET data of cells expressing AKAR4-CAAX in response to 100 nM leptin (n=7), NMDAR co-agonists NMDA/glycine (100 μM/100 μM) (n=8), and AMPK activator AICAR (500 μM) (n=10) in the presence of st-Ht31. Results are compared to those shown in Fig. 2B of cells treated with vehicle (n=13), leptin (n=24), NMDA/glycine (n=11) and AICAR (n=20) in the absence of st-Ht31. *p<0.05, **p<0.01, ***p<0.001 by one-way ANOVA followed by a post-hoc Tukey’s multiple comparison test. B) INS-1 832/13 cells transduced with bungarotoxin binding motif-tagged SUR1 (BTX-SUR1) and Kir6.2 subunits of K_ATP_ channels and treated with leptin (10 nM) for 30 min in the presence of 0.01% DMSO or 50 μM st-Ht31. Surface K_ATP_ channels were then labeled with Alexa 555-conjugated bungarotoxin (BTX) and imaged by confocal microscopy. Nuclei were stained with DAPI. *Scale bar, 10 μm*. C) Representative INS-1 832/13 cell-attached membrane recordings in response to leptin (10 nM) in the absence or presence of st-Ht31 (50 μM) pre-incubation (15-30 min). D) Group data showing the extent of membrane hyperpolarization in response to leptin without st-Ht31 pre-incubation (−64.76 ± 1.61 mV, n=5) or with st-Ht31 pre-incubation (−14.53 ± 3.96 mV, n=8). ***p<0.001 by unpaired student’s t-test. For graph analysis of membrane potential recordings the mean is represented by a thick line with error bars depicting the standard error of the mean.

Having established that PKA-AKAP interactions are necessary for leptin activation of PKA, we wanted to determine if such an interaction is essential for downstream K_ATP_ channel trafficking. To visualize K_ATP_ channels at the plasma membrane, INS-1 832/13 cells were transduced with recombinant adenoviruses containing the K_ATP_ channel subunits Kir6.2 and SUR1 tagged with an N-terminus extracellular bungarotoxin-binding motif (BTX-SUR1)^7^; surface KATP channels were then labeled with Alexa 555-conjugated BTX (BTX) following a 30 min treatment period. Leptin (10 nM) treated cells showed a marked increase in surface BTX staining of BTX-SUR1 compared to vehicle treated cells as expected (Fig. 3B)^7^. However, this effect of leptin was greatly attenuated by the presence of st-Ht31 indicating that blocking PKA from binding AKAPs prohibits leptin from increasing KATP channel surface density. Previously we have shown that the increased abundance of KATP channels in the membrane enhances total K^+^ conductance and causes β-cells to hyperpolarize^7–10^. To further verify the importance of PKA-AKAP interactions for K_ATP_ channel translocation we monitored the effects of st-Ht31 on cell membrane potential following leptin treatment using cell-attached current clamp recording, which is a non-invasive approach for detecting changes in membrane potential without disturbing cellular integrity or compromising intracellular soluble factors important for signaling^34,35^. Cell-attached membrane recordings of INS-1 832/13 cells showed that leptin (10 nM) elicits a mean hyperpolarization extent of −64.76 ± 1.61 mV (n=5) (Fig. 3C, D). However, in the presence of st-Ht31 (50 μM) leptin-induced hyperpolarization was significantly reduced to −14.53 ± 3.96 mV (p< 0.001, n=8). This data strongly implicates that an AKAP is required to anchor PKA for leptin-mediated KATP channel trafficking.

### PKA and AKAPs are also required for leptin signaling in human β-cells

Our previous studies have shown that leptin regulation of K_ATP_ channel trafficking is conserved in human β-cells. To test whether PKA and AKAPs are also involved in leptin signaling in human β-cells, we conducted cell-attached current-clamp recording experiments to monitor cell membrane potential response while pharmacologically manipulating PKA activity and the PKA-AKAP interaction as described above for INS-1 832/13 cells. Individual human β-cells dissociated from islets from three different donors (Table 1) were tested and the results were compared to those using INS-1 832/13 cells (Fig. 4). Bath application of 10 nM leptin induced membrane hyperpolarization in human β-cells to a similar extent (−39.40 ± 6.46 mV, n=11) as in INS-1 832/13 cells (−46.49 ± 4.92 mV, n=11) (Fig. 4A, B). Consistent with PKA being required for KATP channel trafficking, the PKA inhibitor PKI (1 μM) significantly reduced leptin-induced hyperpolarization to −5.18 ± 2.65 mV (p<0.01, n=6) and −17.82 ± 6.88 mV (p<0.01, n=9) in human β-cells and INS-1 832/13 cells, respectively (Fig. 4A, B). Conversely, treatment with the PKA-specific agonist 6-Bnz-cAMP (10 μM) recapitulated the effects of leptin and caused a mean hyperpolarization of −54.84 ± 6.83 mV (n=9) in human β-cells and −56.00 ± 8.15 mV (n=12) in INS-1 832/13 cells. Interestingly, at the concentration used, 6-Bnz-cAMP tended to induce a greater membrane hyperpolarization compared to leptin, which may be due to variation in the degree of PKA activation. The findings in human β-cells agree with those in INS-1 832/13 cells indicating that PKA is both necessary and sufficient to promote K_ATP_ channel trafficking and subsequent membrane hyperpolarization in human β-cells. We then examined whether disrupting the PKA-AKAP interaction would negatively impact leptin-induced hyperpolarization in human β-cells. Pre-incubating human β-cells with st-Ht31 (50 μM) to disrupt PKA-AKAP interactions occluded the effects of leptin on β-cell membrane potential (−4.81 ± 1.13 mV, p<0.0001, n=11). These studies demonstrate the importance of our findings to human biology and support that a PKA-AKAP complex plays an essential role for leptin signaling in human β-cells. In the studies that follow, we will further elucidate the mechanism of PKA-AKAP interaction in leptin signaling using INS-1 832/13 cells as they are more amenable to biochemical and molecular genetic manipulations.

**Figure 4.**
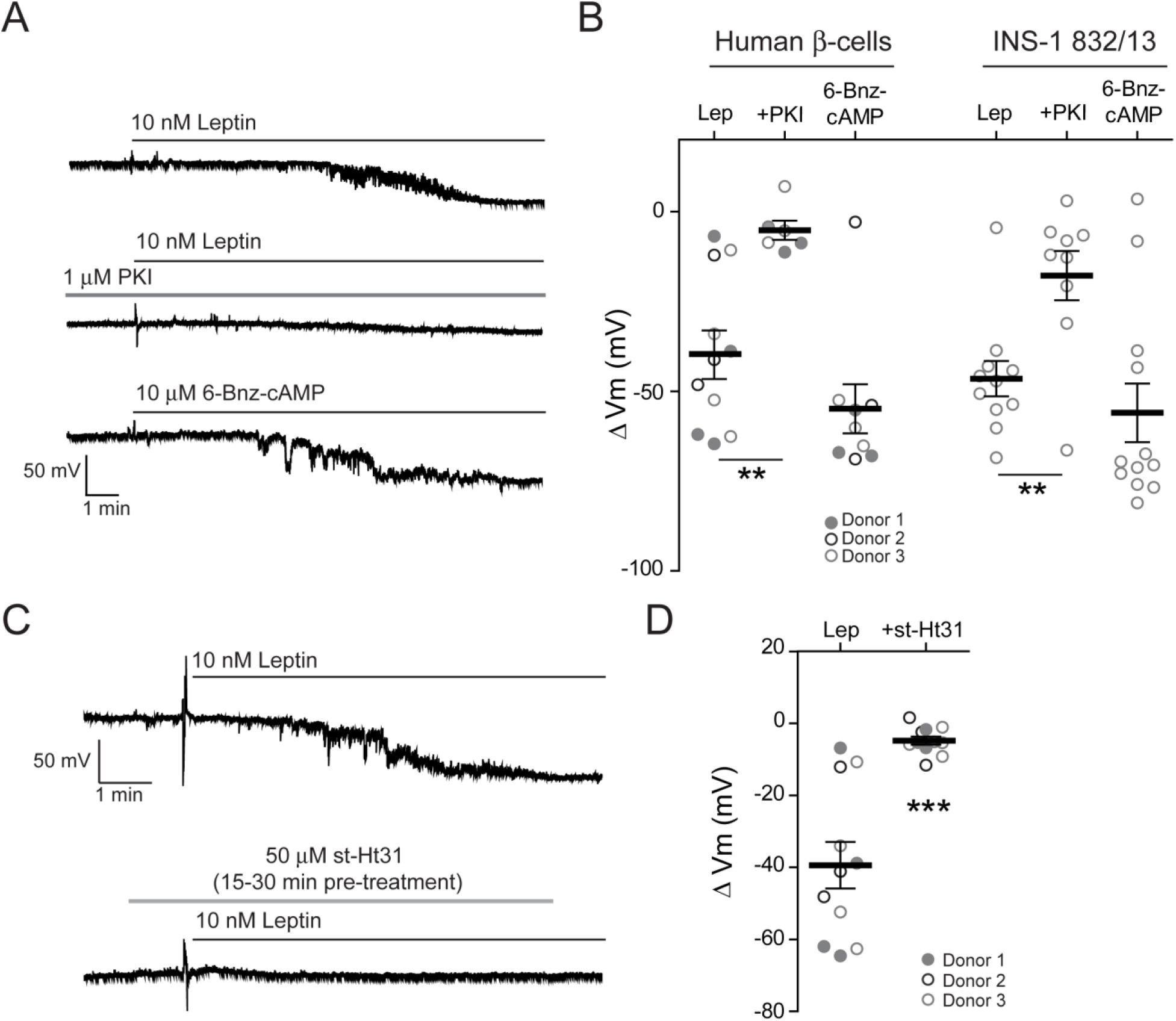
AKAP anchoring of PKA is necessary for leptin-induced hyperpolarization in human β-cells. A) Representative cell-attached membrane recordings of individual human β-cells treated with 10 nM leptin (top), leptin in the presence of the PKA inhibitor PKI (middle; PKI, 1 μM), or the PKA-specific activator 6-Bnz-cAMP (bottom; 6-Bnz-cAMP, 10 μM). B) Group analysis of the extent of membrane hyperpolarization of human β-cells or INS-1 832/13 cells treated with leptin (human β-cells: −39.40 ± 6.46 mV, n=11; INS-1 832/13 cells: −46.49 ± 4.92 mV, n=11), leptin with PKI (human β-cells: −5.18 ± 2.65 mV, n=6; INS-1 832/13 cells: −17.82 ± 6.88 mV, n=9), or 6-Bnz-cAMP (human β-cells: −54.84 ± 6.83 mV, n=9; INS-1 832/13 cells: −56.00 ± 8.15 mV, n=12). **p<0.01 by one-way ANOVA followed by a post-hoc Dunnett’s multiple comparison test. C) Representative membrane potential recordings of human β-cells in response to leptin (10 nM) following pre-incubation without (top) or with 50 μM st-Ht31 (bottom) for 15-30 min. D) Group data of human β-cells showing the degree of membrane hyperpolarization in response to leptin without st-Ht31 pre-incubation (−39.40 ± 6.46 mV, n=11) or with st-Ht31 pre-incubation (−4.81 ± 1.13 mV, n=11). ***p<0.0001 by unpaired student’s t-test. Donors are indicated by the circle color and fill.

**Table 1.**
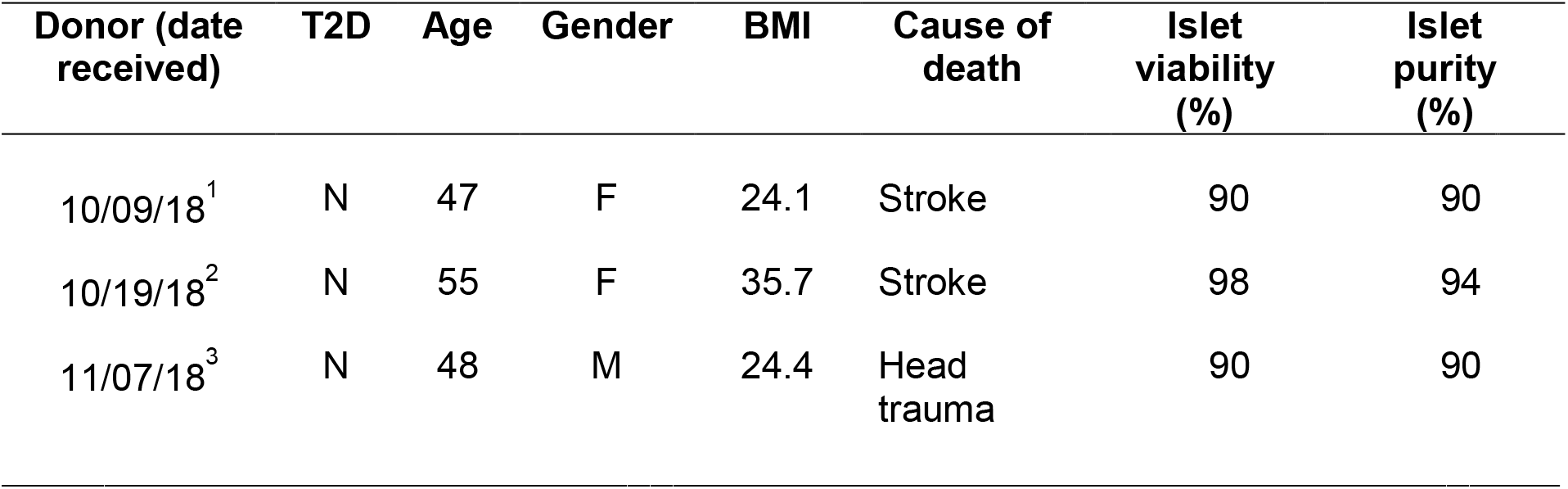
Human Islets Donor Information.

### AKAP79/150 coordinates leptin signaling to regulate K_ATP_ channel trafficking

The above results that PKA-AKAP interactions are required for leptin to exert its signaling effect raises the question of which AKAP(s) is involved in localizing PKA. There are more than 50 known AKAPs^19^. We chose to focus on AKAPs that have been shown to be expressed in β-cells and localize at the cell membrane based on the observation that PKA activation occurs near the plasma membrane upon leptin stimulation (Fig.1). Among the possible candidates, AKAP79/150 (human/murine proteins, *AKAP5* gene) stood out as a most interesting candidate^20,36–38^, because in neurons AKAP79/150 has been shown to co-immunoprecipitate with NMDARs^39,40^, a known player in the leptin signaling pathway being studied here. To test the idea that AKAP79/150 could serve as a scaffold to coordinate a complex of leptin signaling molecules in β-cells and regulate K_ATP_ channel trafficking, we genetically knocked down AKAP150 expression and monitored surface K_ATP_ channels using surface staining, electrophysiology and surface biotinylation experiments.

INS-1 832/13 cells were transiently transfected with AKAP150 shRNAi to knock down AKAP150 (AKAP150 KD) or with the empty pSilencer vector as a control ^41,42^. AKAP150 KD cells showed a significant decrease in AKAP150 expression compared to control cells regardless of whether they received vehicle or leptin (10 nM) treatment for 30 minutes (Fig. 5A). Surface staining of exogenously expressed BTX-SUR1/Kir6.2 K_ATP_ channels was again used to visualize K_ATP_ channels in the membrane. Similar to untransfected INS-1 832/13 cells (Fig. 3B), pSilencer transfected control cells also showed increased surface K_ATP_ channel expression when treated with leptin, but this effect was abolished in AKAP150 KD cells (Fig. 5B). In agreement with this finding, cell-attached current-clamp recordings showed that leptin-induced hyperpolarization was drastically reduced from −60.40 ± 4.60 mV (n=5) in pSilencer control cells to 0.30 ± 0.49 mV (p<0.0001, n=6) in AKAP150 KD cells (Fig. 5C). To further substantiate the role of AKAP150 in leptin signaling we carried out surface biotinylation experiments to examine the expression of endogenous K_ATP_ channels at the membrane. Control and AKAP150 KD cells were treated with vehicle or leptin (10 nM) for 30 minutes followed by surface biotinylation and western blot analysis. In pSilencer control cells leptin caused a 2.39-fold increase (p<0.05, n=3) in surface SUR1 compared to vehicle treatment (Fig. 5D). In contrast, AKAP150 KD cells failed to show an increase in surface K_ATP_ channels upon leptin treatment. Taken together these findings identify a novel relationship between AKAP79/150 and leptin signaling in β-cells.

**Figure 5.**
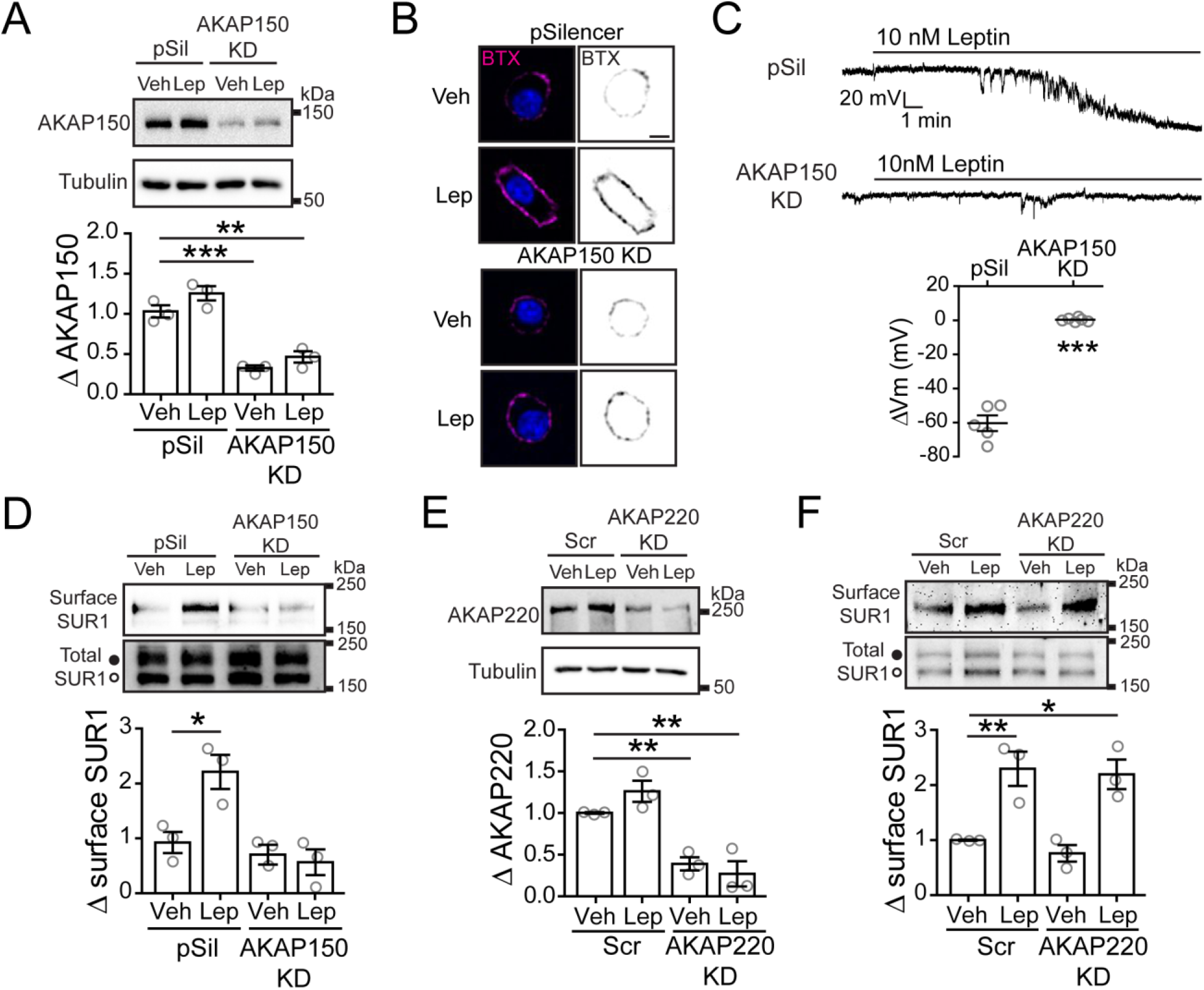
AKAP150 is necessary for leptin-induced K_ATP_ channel trafficking. A) INS-1 832/13 cells transfected with the control pSilencer vector (pSil) or AKAP150 shRNAi (AKAP150 KD) and analyzed for AKAP150 expression via western blot (top). Graph shows quantification of AKAP150 relative to tubulin (bottom). B) Confocal images of BTX labeled surface KATP channels following vehicle or 10 nM leptin treatment in pSil and AKAP150 KD cells. *Scale bar, 5 μm*. C) Representative membrane potential recordings from pSil (top; −60.40 ± 4.60 mV, n=5) and AKAP150 KD (bottom; 0.30 ± 0.49 mV, n=6) cells treated with 10 nM leptin. Below the traces is the group analysis of the extent of membrane hyperpolarization. ***p<0.0001 by unpaired student’s t-test. D) Surface biotinylation experiments. Western blots show surface expression of the K_ATP_ channel subunit SUR1 and total SUR1 in pSil and AKAP150 KD cells treated with vehicle or 10 nM leptin (top). Note, the upper band in the total SUR1 corresponds to the complex-glycosylated SUR1 (filled circles) that traffics to the surface and the lower band corresponds to the ER-core glycosylated SUR1 (open circle). Quantification of surface SUR1 relative to total SUR1 (bottom). E) Western blot analysis (top) and quantification (bottom) of AKAP220 expression in control scramble siRNA (Scr) or AKAP220 siRNA (AKAP220 KD) cells. F) Western blots showing the effects of AKAP220 siRNA on surface SUR1 relative to total SUR1. Biochemical experiments (A, D, E, F) were repeated three times (n=3; represented as circles) and normalized to vehicle treated controls. *p<0.05, **p<0.01, ***p<0.001 by one-way ANOVA followed by a post-hoc Dunnett’s multiple comparison test unless stated otherwise.

To determine whether the effect observed with AKAP150 KD is specific, we examined the effect of knockdown of another membrane associated AKAP, AKAP220^43,44^. INS-1 832/13 cells were transfected with scramble control siRNA or AKAP220 siRNA (AKAP220 KD) and knockdown of AKAP220 expression was confirmed by western blot analysis (Fig. 5E). Surface biotinylation experiments were then carried out to assess K_ATP_ channel trafficking in response to leptin. As expected, scramble control cells treated with leptin showed a 2.30-fold increase (p<0.01, n=3) in surface SUR1 expression compared to vehicle control (Fig. 5F). AKAP220 KD cells stimulated with leptin also showed a significant 2.19-fold increase (p<0.01, n=3) in surface SUR1 expression compared to vehicle treated scramble control cells. The amount of surface SUR1 in leptin treated AKAP220 KD cells was comparable to scramble control cells treated with leptin and in stark contrast to AKAP150 KD cells. From these observations we concluded that AKAP220 is not involved and that AKAP79/150 plays a unique role for leptin signaling.

### AKAP79/150 mediates leptin signaling via its interaction with PKA

As a PKA anchoring protein, AKAP79/150 presumably participates in leptin signaling by binding to PKA and bringing PKA in proximity to the other signaling molecules. However, in addition to PKA, AKAP79/150 has also been found to anchor protein phosphatase 2B (PP2B; also known as calcineurin; see Fig. 6A)^45,46^, which opposes PKA activity by dephosphorylating PKA substrates. This dual specificity of AKAP79/150 coordinates PKA and PP2B signaling and allows anchored-PKA actions on downstream substrates to be tightly regulated. Indeed, AKAP79/150 mutants that lack the PP2B binding PxIxIT-like motif (AKAP79ΔPIX; Fig. 6A)^45–47^ display increased localized PKA activity^42,45,47,48^. To address whether the role of AKAP79/150 in leptin signaling involves AKAP79/150 interaction with PKA, PP2B, or both, we performed AKAP150 KD and rescue experiments using full-length WT AKAP79 (human ortholog of mouse AKAP150), or AKAP79 mutants deficient in PKA or PP2B binding.

**Figure 6.**
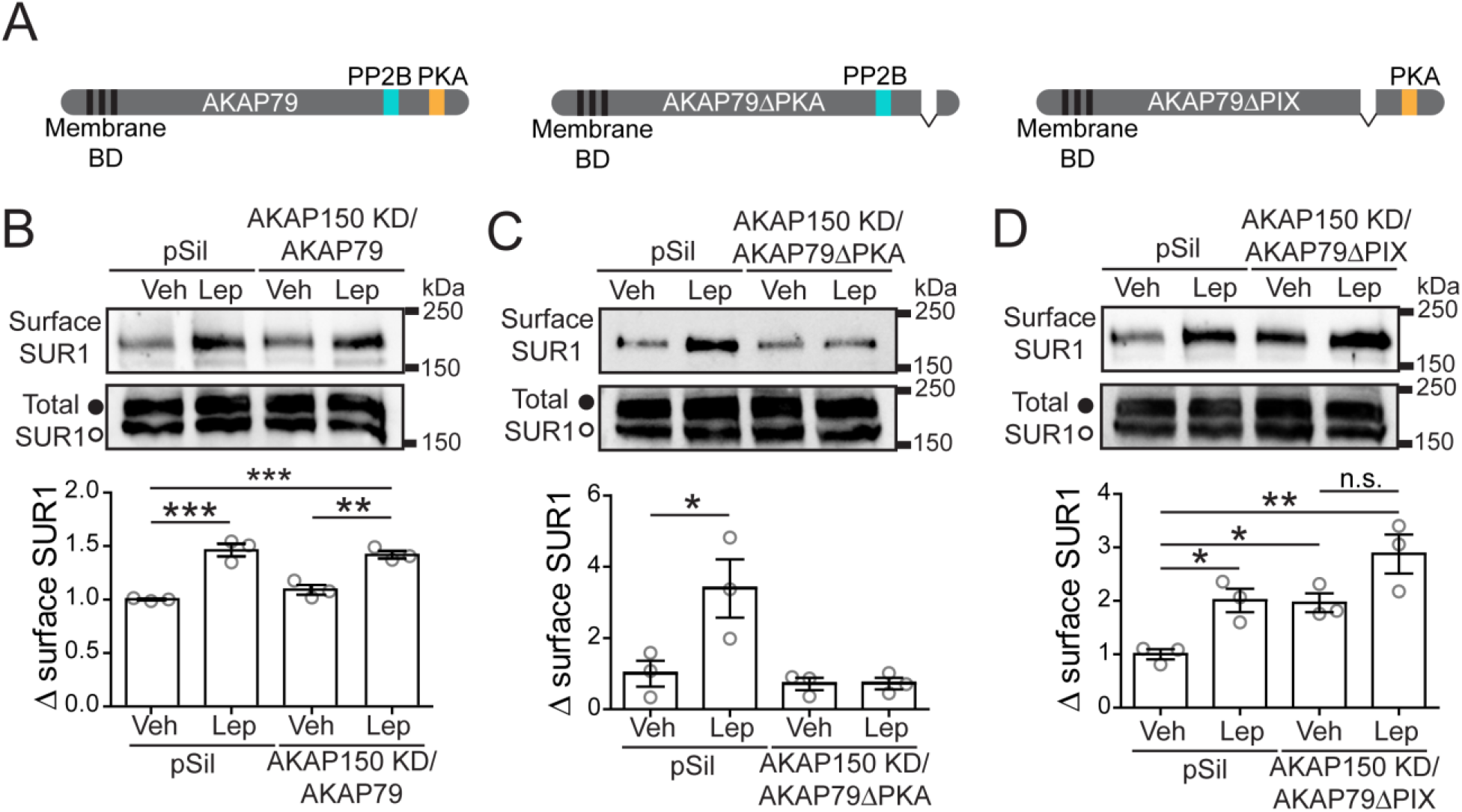
AKAP79 rescues leptin-induced K_ATP_ channel trafficking in AKAP150 KD cells. A) Schematic of AKAP79 showing key membrane binding domains (BD) as well as PP2B and PKA binding regions. B) INS-1 832/13 cells were transfected with control pSilencer vector (pSil) or co-transfected with AKAP150 shRNAi and WT AKAP79 (AKAP150 KD/AKAP79). Transfected cells were treated with vehicle or 10 nM leptin for 30 min followed by surface biotinylation. Western blots show surface SUR1 and total SUR1 (top). Quantification of surface SUR1 relative to total SUR1 (bottom). C, D) Same as (B) except AKAP150 KD cells were co-transfected with AKAP79 mutants that cannot bind PKA (AKAP150 KD/AKAP79ΔPKA) or PP2B (AKAP150 KD/AKAP79ΔPIX). Each experiment was performed three independent times (n=3; shown as circles) and results were normalized to vehicle treated control pSil cells. *p<0.05, **p<0.01, ***p<0.001 by one-way ANOVA followed by a post-hoc Tukey’s multiple comparison test.

First, we performed surface biotinylation experiments in AKAP150 KD cells transfected with WT AKAP79 (AKAP150 KD/AKAP79) to test whether expression of AKAP79 would rescue leptin-induced K_ATP_ channel trafficking in AKAP150 KD cells. In this set of experiments leptin caused a 1.47-fold increase (p<0.001, n=3) of surface SUR1 in pSilencer transfected control cells. A similar 1.42-fold increase (p<0.001, n=3) was observed in AKAP150 KD/AKAP79 cells demonstrating that AKAP79 successfully rescues a loss of AKAP150 (Fig. 6A, B). This result also confirms that the lack of leptin response in AKAP150 KD cells (Fig. 5D) was not due to off-target effects of the shRNAi. Next, we performed the same experiment using an AKAP79 mutant that lacks the PKA binding motif (AKAP79ΔPKA; Fig. 6A)^46,47^ to test the importance of AKAP79/150-anchored PKA for leptin signaling. AKAP150 KD/AKAP79ΔPKA cells treated with leptin failed to show a significant change in surface KATP channel expression compared to vehicle treated cells (Fig. 6C) indicating that PKA anchored by AKAP79/150 serves a critical function in the leptin signaling cascade. Lastly, surface biotinylation experiments were performed in AKAP150 KD cells transfected with an AKAP79 mutant lacking the PP2B binding motif (AKAP150 KD/AKA79ΔPIX). Interestingly, compared to vehicle controls, vehicle treated AKAP150 KD/AKA79ΔPIX cells exhibited a 1.96-fold increase (p<0.05, n=3) in surface SUR1 expression levels akin to that seen in leptin treated controls (2.01-fold increase, p<0.05, n=3) (Fig. 6D). Stimulating AKAP150 KD/AKA79ΔPIX cells with leptin led to an even greater 2.77-fold increase (p<0.01, n=3) in surface K_ATP_ channels, although this increase was not statistically significant when compared to vehicle treated AKAP150 KD/AKA79ΔPIX cells (p>0.05). The higher than normal K_ATP_ channel surface expression in unstimulated AKAP150 KD/AKAP79ΔPIX cells implies that PP2B constitutively bound to AKAP79 likely limits anchored-PKA signaling such that disrupting PP2B anchoring mimics the effect of leptin to increase basal PKA signaling and K_ATP_ channel trafficking. By increasing basal PKA signaling AKAP79ΔPIX could partially mask the effect of leptin on K_ATP_ channel trafficking, thus explaining the lack of a statistically significant difference between leptin and vehicle treated groups. The combined results from these experiments provide compelling evidence that leptin activates AKAP79/150-anchored PKA and that these actions of PKA are basally opposed by AKAP79/150-anchored PP2B.

### Leptin increases cAMP concentrations near AKAP79

The results thus far indicate that leptin increases the activity of AKAP79/150 anchored-PKA to regulate K_ATP_ channel trafficking, but how leptin activates PKA remains to be addressed. PKA is typically activated by cAMP, which binds to PKA regulatory subunits and relieves autoinhibition to unleash the catalytic subunits. Alternatively, inhibiting the opposing activity of PP2B could also enhance net PKA signaling^11,31,47–49^. Our results above implicating that AKAP150 KD/AKAP79ΔPIX cells have increased basal PKA signaling is consistent with an inhibitory role of PP2B on steady state PKA signaling. However, the observation that AKAP150 KD/AKAP79ΔPIX cells still responded to leptin suggests a PKA activation mechanism independent of PP2B inhibition. This prompted us to ask whether leptin increases cAMP levels near AKAP79/150. To monitor changes in cAMP near AKAP79/150, we employed the FRET-based cAMP CUTie sensor targeted to AKAP79 (AKAP79-CUTie)^50^. AKAP79-CUTie was found to express at the cell membrane where we had also observed increases in PKA activity. Similar to FRET-based PKA activity experiments, we treated the cells with vehicle or leptin (100 nM), normalized the FRET traces to a maximal response, and then analyzed the traces for area under the curve. To generate a maximal response we applied forskolin (20 μM) and IBMX (10 μM), which increases cAMP production by adenylyl cyclases (ACs) and prevents cAMP degradation by phosphodiesterases (PDEs), respectively. During our first set of experiments we did not see a significant difference in cAMP levels between treatments, although there appeared to be a trend toward increased cAMP in leptin treated cells (Fig. 7A). Since AKAPs have also been shown to anchor PDEs to tightly regulate PKA activation via cAMP degradation^19^ we repeated the experiments in the presence of IBMX to prevent potential cAMP degradation that could obscure cAMP signals. In the presence of IBMX, leptin treated cells showed a significant 1.53-fold increase (p<0.0005, n=16) in AKAP79-CUTie cAMP sensor activity compared to vehicle (n=15) (Fig. 7B). This effect of leptin on cAMP levels appears to be specific to AKAP79 as leptin did not significantly increase cAMP levels monitored by a cytoplasmic AKAP18δ-CUTie sensor (Fig. 7C). These results lead us to conclude that leptin signaling activates PKA at least in part by increasing local cAMP levels near AKAP79/150.

**Figure 7.**
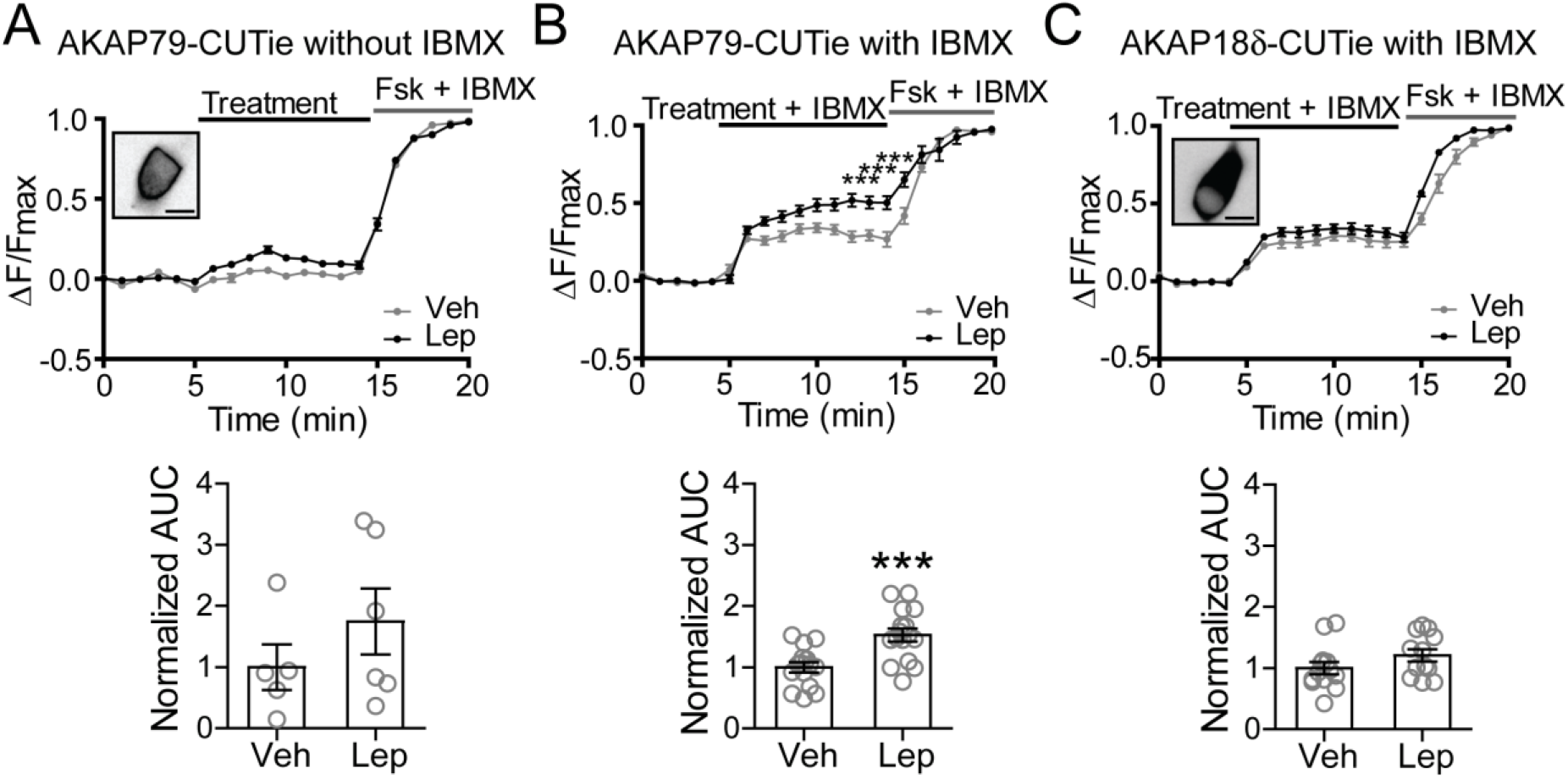
Leptin increases cAMP levels near AKAP79. A) cAMP CUTie sensor targeted to AKAP79 (human orthologue of AKAP150) was utilized to detect changes in cAMP levels in response to vehicle or 100 nM leptin. The average FRET traces (top) and the group data of FRET traces analyzed for area under the curve (bottom) are shown. *Inset shows INS-1 832/13 cell expressing AKAP79-CUTie sensor. Scale bar, 10 μm*. B) Experiments in (A) were repeated in the presence of the phosphodiesterase inhibitor IBMX (50 μM) to prevent rapid degradation of cAMP. ***p<0.0005 as determined by unpaired student’s t-test. C) Same as (B) except cAMP levels were monitored using the AKAP18δ-CUTie sensor, which is largely expressed in the cytosol as shown in the inset. *Scale bar, 10 μm*.

## Discussion

The study we present here reveals a novel relationship between leptin and PKA for K_ATP_ channel trafficking in β-cells that is mediated by AKAP79/150. From this work and our previous studies we propose a model in which the PKA-AKAP79/150 complex orchestrates leptin-induced K_ATP_ channel trafficking to suppress GSIS from pancreatic β-cells (Fig. 8). In this model leptin stimulation increases PKA activity anchored by AKAP79/150 through a signaling cascade wherein leptin activates Src kinase which phosphorylates and potentiates NMDARs, resulting in an enhanced Ca^2+^ influx that activates CaMKKβ to phosphorylate and activate AMPK. AMPK then increases PKA activity by elevating cAMP concentrations localized near AKAP79/150, possibly by activating an AC, culminating in actin remodeling and the translocation of K_ATP_ channels. A greater abundance of K_ATP_ channels in the membrane increases the total K^+^ conductance causing β-cell membrane hyperpolarization and inhibition of Ca^2+^ influx through voltage-dependent Ca^2+^ channels to prevent insulin exocytosis. Importantly, we found that in human β-cells the ability of leptin to reduce electrical activity was also dependent on PKA-AKAP interactions suggesting that this signaling mechanism serves an important function in the regulation of insulin secretion in humans.

**Figure 8.**
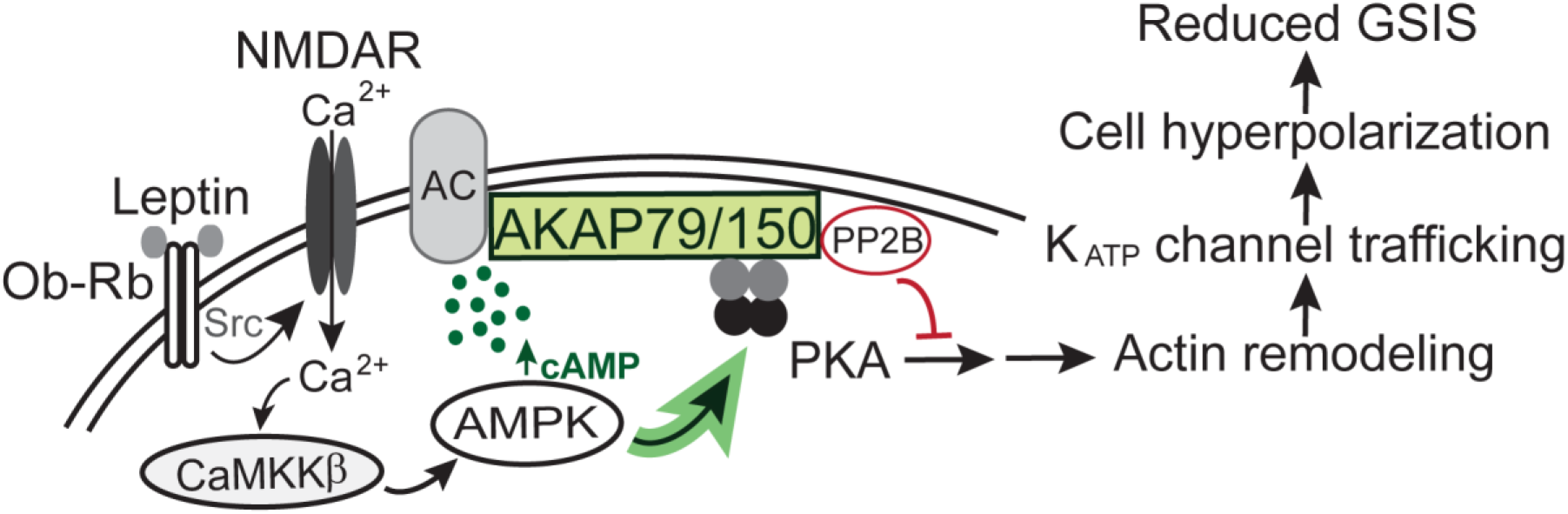
Proposed model depicting AKAP79/150 mediates leptin signaling to regulate K_ATP_ channel trafficking. AKAP79/150 is a scaffolding protein that creates PKA signaling microdomains localized at cell membranes. AKAP79/150 also anchors PP2B, which opposes PKA activity by dephosphorylating PKA substrates and ACs, which enhance PKA activity by producing the PKA activator cAMP. Signaling complexes coordinated by AKAP79/150 allow for PKA signaling to be tightly regulated. In pancreatic β-cells AKAP79/150 anchoring of PKA renders a localized increase of PKA activity following leptin activation of Src kinase to initiate the NMDAR-CaMKKβ-AMPK signaling cascade. This activation of PKA may at least in part be due to the leptin signaling axis increasing cAMP levels near AKAP79/150. Actin remodeling downstream of PKA activation allows for increased KATP channel trafficking and a subsequent increase in K^+^ conductance, which causes cell hyperpolarization and suppresses GSIS.

### Mechanism of leptin activation of PKA

Although our previous studies suggested that PKA-dependent actin remodeling was necessary for leptin to promote K_ATP_ channel trafficking^7^, there was very little evidence outside of our own findings for such a signaling relationship between leptin and PKA^51^. By implementing FRET-based live cell imaging to monitor PKA activity in combination with pharmacology we demonstrate here that leptin significantly increases PKA activity at the cell membrane via the NMDAR-CaMKKβ-AMPK signaling axis. The finding that AMPK activation can increase PKA activity in β-cells was notable as two prior studies, one in vascular smooth muscle cells and the other in cardiomyocytes, had only tentatively implicated AMPK upstream of PKA^52,53^. One potential mechanism for AMPK to increase PKA activity is by inhibiting protein phosphatase activity to prevent the dephosphorylation of PKA effectors. While there are some studies to suggest that AMPK may counteract protein phosphatases^54^, many instead show that AMPK is inactivated by protein phosphatases^55,56^ or that AMPK increases protein phosphatase activity^57,58^. Moreover, our experiments using the AKAP79ΔPIX mutant that cannot bind PP2B indicates that leptin signaling may increase PKA activity independent of PP2B. Alternatively, our finding that leptin stimulation raises cAMP levels near AKAP79-CUTie suggests that this may be how AMPK increases PKA activity. It is unlikely that AMPK increases cAMP by inhibiting PDEs as we were only able to detect significantly elevated cAMP levels in the presence of the PDE inhibitor IBMX, and in other contexts AMPK has been shown to increase PDE activity^59^. Rather, AMPK may act via ACs either directly or indirectly to cause a rise in cAMP. In this regard, it is interesting to note that several ACs have been reported to interact with AKAP79/150^21,60–64^ Other possibilities that AMPK phosphorylates a yet-to-be identified intermediary or directly phosphorylates PKA to induce catalytic activity also need to be considered^65^. It is clear that more work is required to determine the precise mechanism by which AMPK activates PKA.

### Spatiotemporal regulation of PKA activity in β-cells

A key observation made during these studies is that leptin causes an increase in PKA activity that is restricted near the cell membrane. While our studies support that leptin signals via PKA to increase K_ATP_ channel surface density, K^+^ conductance and subsequent β-cell hyperpolarization known to inhibit GSIS^7,8^, others have found that the incretin hormone glucagon-like peptide 1 (GLP-1) signals via PKA to promote insulin granule trafficking^14^ and Ca^2+^-induced exocytosis to augment GSIS^15–17,23^. All of these studies have used pharmacological or genetic manipulations of total PKA activity to implicate PKA in either inhibiting or enhancing GSIS. Although some of these discrepancies may be explained in part by differences in experimental design, it is also likely that they are the result of the indiscriminate activation and/or inhibition of total cellular PKA. In contrast, endogenous hormonal and physiological stimuli such as leptin and GLP-1 are likely to act in a more nuanced manner due to high spatial and temporal regulation. This is supported by our data demonstrating that leptin has a much more limited effect on PKA activity compared to the global PKA activator forskolin. Specificity of PKA signaling within cells is attributed primarily to AKAPs, such as AKAP79/150, organizing PKA signaling microdomains. Interestingly, recent work in vascular smooth muscle cells also illustrates how local cAMP-PKA signaling in an AKAP79/150- organized complex can dictate a very different downstream cellular response from that regulated by more global engagement of cAMP and PKA^66^. Of note, some AKAPs expressed in β-cells, including AKAP79/150, have already been implicated in either inhibiting or promoting insulin secretion^20–25^. This highlights that a high level of structural cellular organization is essential for β-cells to incorporate multiple complex signaling networks in order to function properly, including the opposing effects of the hormones leptin and GLP-1.

### Implications for the role of AKAP79/150 in insulin secretion

Our results that AKAP150 knockdown or expression of AKAP79ΔPKA prevents leptin-induced K_ATP_ channel trafficking indicate that AKAP79/150 coordinates PKA and leptin signaling to impact insulin secretion. Interestingly, neither AKAP150 knockdown nor AKAP79ΔPKA expression affected basal K_ATP_ channel surface expression, suggesting that while AKAP79/150 anchored-PKA is critical for leptin signaling it is unlikely to be involved in constitutive K_ATP_ channel trafficking. On the other hand, our findings that expression of AKAP79ΔPIX, a mutant that cannot bind PP2B, leads to a greater abundance of K_ATP_ channels in the β-cell membrane in the absence of leptin stimulation suggests that increasing AKAP79/150 anchored-PKA activity is sufficient to promote K_ATP_ channel translocation. Note, a previous study has reported that mouse islets lacking AKAP150 exhibit reduced GSIS^20^, which was attributed to the effects of AKAP150 on L-type Ca^2+^ channels. Curiously, the same study found that while expressing AKAP150ΔPKA in AKAP150 knockout islets did not affect GSIS, expressing AKAP150ΔPIX significantly reduced GSIS^20^. These observations are congruent with our findings in INS-1 832/13 cells that disrupting PKA tethering did not affect basal surface K_ATP_ channel density but disrupting PP2B tethering increases basal surface K_ATP_ channel density. This raises the possibility that the reduced GSIS observed in AKAP150ΔPIX-expressing islets is in part due to increased K_ATP_ channel surface expression. Also important, while our current study focuses on the role of AKAP79/150 in mediating PKA activation by leptin, AKAP79/150 is multifaceted and can coordinate multiple signaling complexes, as has been shown in neurons^39,42,67^ and other cell types^66,68^. Thus, AKAP79/150 in β-cells may be involved in organizing multiple signaling networks with distinct functions. Finally, increased K_ATP_ channel trafficking has been observed following glucose starvation via activation of AMPK^69^, and following high glucose stimulation in a PKA-dependent manner^70^. It will be interesting to determine in the future whether these regulations are also mediated by AKAP79/150.

In summary, we have identified a novel role of AKAP79/150 in coordinating leptin and PKA signaling to regulate K_ATP_ channel trafficking in β-cells, hence GSIS. Importantly, our recent studies showed that in human β-cells from obese type II diabetic donors and β-cells from obese diabetic *db/db* mice lacking functional leptin receptors leptin fails to promote K_ATP_ channel trafficking and membrane hyperpolarization; however, activation of NMDARs downstream of leptin reenacted the effect of leptin^10^. Thus further exploration into the leptin signaling pathway coordinated by AKAP79/150 will likely provide valuable insight into how the β-cell regulates K_ATP_ channel trafficking to tune its function and may even identify potential therapeutic targets to combat type II diabetes.

## Methods

### Chemicals

Leptin and glutamate were from Sigma-Aldrich (St. Louis, MO). PKI, 6-Bnz-cAMP, NMDA, Compound C (Dorsomorphin), D-APV, and STO-609 were from Tocris Bioscience (Bristol, UK). AICAR was from Selleck Chemicals (Houston, TX). st-Ht31 inhibitor peptide was from Promega (Madison, WI).

### INS-1 832/13 cell culture

INS-1 cells (clone 832/13, referred to herein as INS-1 832/13) were cultured in RPMI 1640 medium with 11.1 mM D-glucose (Invitrogen) supplemented with 10% fetal bovine serum (FBS), 100 units/ml penicillin, 100 μg/ml streptomycin, 10 mM HEPES, 2 mM glutamine, 1 mM sodium pyruvate, and 50 μM β-mercaptoethanol. Only cells within passage numbers 55-75 were used for experiments.

### Dissociation of human pancreatic β-cells

Human β-cells were dissociated from human islets obtained through the Integrated Islets Distribution Program (IIDP) as described previously^7–10^. Human islets were cultured in RPMI 1640 medium with 10% FBS and 1% L-glutamine. Islets were dissociated into single cells by trituration in a solution containing 116 mM NaCl, 5.5 mM D-glucose, 3 mM EGTA, and 0.1% bovine serum albumin (BSA), pH 7.4. Dissociated cells were then plated on 0.1% gelatin-coated coverslips and allowed to recover overnight in culture media. For electrophysiological experiments, β-cells were identified by their high level of autofluorescence at 488 nm excitation due to β-cells having high concentrations of unbound flavin adenine dinucleotide^71,72^. Dithizone (Sigma-Aldrich) staining confirmed β-cell identity at the end of each experiment^73^. Donor information is provided in Table 1.

### Plasmids, viruses, and siRNA

pcDNA3-EGFP (Addgene plasmid #13031) was a gift from Doug Golenbock (UMass, MA). pcDNA3-AKAR4-CAAX (Addgene plasmid #61621) and pcDNA3-AKAR4-NES (Addgene plasmid #64727) were gifts from Jin Zhang (UCSD, CA). AKAP79-CUTie and AKAP18δ-CUTie were gifts from Manuela Zaccolo (Oxford, UK). AKAP150 shRNAi and pSilencer vector were gifts from John Scott (UW, Washington)^41^, AKAP79-GFP, AKAP79ΔPIX, and AKAP79ΔPKA were described previously^36,42,45,47^. Recombinant rat Kir6.2 and BTX-tag SUR1 adenoviruses were generated in our lab and described previously^7^. AKAP220 siRNA (5’-CCAAUGUAAGCA GUAGUCCUCUAA A-3’) and scramble siRNA (5’-UUUAGAGGACUACUGCUUACA UUGG-3’) were from Millipore (Burlington, MA).

### Electrophysiology

Cell-attached recordings were performed using an Axon 200B amplifier (Molecular Devices, Sunnyvale, CA) and pClamp software. Signals were acquired at 20 kHz and filtered at 2 kHz. Micropipettes were pulled from non-heparinized Kimble glass (Thermo Fisher Scientific, Waltham, MA) on a horizontal puller (Sutter Instruments, Novato, CA) and filled with 140 mM NaCl. The bath solution (Tyrode’s solution) contained (in mM): 137 NaCl, 5.4 KCl, 1.8 CaCl_2_, 0.5 MgCl_2_, 5 Na-HEPES, 3 NaHCO_3_, and 0.16 NaH_2_PO_4_, 11 glucose, pH 7.2. Pipette resistance was typically between 2-6 MΩ. The seal resistance between the recording pipette and the cell ranged between 2-8 GΩ. Membrane potentials were recorded in current clamp mode (I=0) and signals were analyzed using Clampfit (pClamp). After correcting for the measured liquid junction potential (−10 mV), the average baseline V_m_ was around −4 mV. In this configuration the estimated Vm is dependent on the ratio of the seal resistance and the combined patch and cell resistance where the ratio of recorded Vm and true membrane potential = (R_seal_/R_patch_+R_cell_)/[1+(R_seal_/R_patch_+R_cell_)]^35^. Note, as the recorded V_m_ is an underestimate of the actual membrane potential we only used it to track changes in membrane potential^34,35^. Seal resistance was monitored before and after the recording. Only cells that showed stable baseline membrane potential prior to leptin/drug application and which maintained good seal resistance were included for analysis.

### Live cell FRET imaging

INS-1 832/13 cells were grown in 35-mm dishes and transfected at 50-60% confluency with FRET-based biosensors (2.5 μg plasmid DNA) using Lipofectamine 2000 (6 μl) (Invitrogen). 24 hours post-transfection cells were plated in μ-slide 8 well glass bottom chamber (Ibidi) and allowed to recover overnight. Cells were then imaged at 37°C in Tyrode’s solution (described above). Images were acquired every minute using an Olympus IX71 inverted microscope with a 40x, 1.35 NA, oil immersion, UApo objective (Olympus). The microscope was equipped with a Nikon Coolsnap ES2 HQ camera. Image processing was performed using Fiji software (National Institutes of Health)^74^. FRET was calculated as the ratio of acceptor fluorophore emission (545 nm) to donor emission (480 nm) in response to donor excitation (435 nm). F/Fmax was obtained for each time point by subtracting the average baseline FRET ratio and normalizing to the maximal forskolin response. Only cells that showed a 10-15% increase in FRET in response to forskolin were used for analysis. Area under the curve (AUC) was calculated using GraphPad Prism™ for each experimental treatment time course.

### Surface staining

INS-1 832/13 cells at ~70% confluency were washed once with phosphate-buffered saline (PBS) and incubated for two hours at 37°C in Opti-MEM (Thermo Fisher Scientific) and a mixture of viruses: tetracycline-inhibited transactivator, tetracycline-inhibited transactivator-regulated construct expressing BTX-SUR1 and Kir6.2^7^. The multiplicity of infection for each virus was determined empirically. After two hours the infection mixture was replaced with RPMI 1640 supplemented cell culture media (described above) and were incubated at 37°C. 24 hours post-infection cells were plated on 15-mm, number 1.5 glass coverslips (Thermo Fisher Scientific) and allowed to adhere overnight. In Fig. 3B cells were preincubated at 37°C for 30 min in RPMI 1640 without serum and an additional 30 min with 0.01% DMSO or 50 μM st-Ht31 before being treated with 10 nM leptin or vehicle for 30 min. A prior study by our lab determined that leptin stimulation for 30 min is the optimal treatment duration for immunocytochemistry and biochemical experiments^7^. Following leptin or vehicle treatment surface BTX-SUR1 was labeled with 1 μg/ml Alexa Fluor^®^ 555 α-bungarotoxin (555-BTX, Thermo Fisher Scientific) for 1 hour at 4°C. In Fig. 5C cells were co-transfected with AKAP150 shRNAi and EGFP or pSilencer and EGFP at a 5:1 plasmid DNA ratio respectively using Lipofectamine 2000. The next day cells were infected with BTX-SUR1 and Kir6.2. These cells were treated with vehicle or 10 nM leptin for 30 min and surface BTX-SUR1 was labeled as described for Fig. 3B (minus the pre-incubation with DMSO or st-Ht31). All imaging experiments were performed on a Zeiss LSM780 confocal microscope equipped with a 63x, 1.4 NA, oil immersion, PlanApochromat objective (Carl Zeiss). During imaging for Fig. 5B EGFP expression was used to distinguish which cells had been transfected. Images were processed with Fiji (NIH).

### Immunoblotting

INS-1 832/13 cells were lysed in triple lysis buffer [50 mM Tris-HCl, 2 mM EDTA, 2 mM EGTA, 100 mM NaCl, 1% Triton X-100, pH 7.4, with cOmplete EDTA free protease inhibitor cocktail (Roche, Basel, Switzerland)] for 30 min at 4°C with rotation, and cell lysates were cleared by centrifugation at 21,000 × *g* for 10 min at 4°C. Proteins were separated by SDS-PAGE (8% acrylamide gel) and transferred to nitrocellulose membranes (Millipore). Membranes were incubated overnight at 4°C with a primary antibody diluted in Tris-buffered saline plus 0.1% Tween 20 (TBST). The antibody for SUR1 (1:500) was generated in rabbit using a C-terminal peptide (KDSVFASFVRADK) of hamster SUR1 as described previously^7^. The antibody against AKAP150 (1:1000) was made as described previously^75^. The antibody against GFP (1:100) was from Thermo Fisher Scientific. The antibody for tubulin (1:2000) was from Sigma-Aldrich. After three 10 min washes with TBST, blots were incubated for 1 hour at room temperature with horseradish peroxidase-conjugated secondary antibodies in TBST buffer as follows: 1:20,000 donkey anti-rabbit IgG (Jackson ImmunoResearch Laboratories) for SUR1, AKAP150, and GFP; 1:20,000 donkey anti-mouse IgG (Jackson ImmunoResearch Laboratories) for AKAP220 and tubulin. After washing three times for 10 min with TBST, blots were developed using Super Signal West Femto (Pierce) and imaged with FluorChemE (ProteinSimple, San Jose, CA) or Sapphire Biomolecular Imager (Azure Biosystems). The blots were quantified using Fiji software (NIH) and normalized to corresponding controls.

### Surface Biotinylation

INS-1 832/13 cells cultured in 10 cm dishes at 50-60% confluency were transfected with 20 μg of pSilencer or AKAP150 shRNAi plasmid DNA and 40 μl of lipofectamine 2000. 48 hours following transfection the cells were re-plated in new 10 cm dishes. 72 hours post-transfection cells at 70-80% confluency were incubated in RPMI 1640 without serum for 1 hour at 37°C prior to a 30 min treatment with vehicle or 10 nM leptin. Cells were then washed four times with cold PBS containing 9 mM CaCl2 and 5.9 mM MgCl2 (DPBS) and incubated with 1 mg/ml EZ-Link Sulfo-NHS-SS-Biotin (Pierce) in DPBS for 30 min at 4°C. The reaction was terminated by incubating cells twice with DPBS containing 50 mM glycine for 5 min at 4°C, followed by two washes with cold DPBS. Cells were then lysed with 300 μl triple lysis buffer as described above, and 500 μg of total lysate was incubated with 50 μl of 50% slurry Neutravidin-agarose beads (Pierce) overnight at 4°C. Biotinylated proteins were eluted with 2x protein loading buffer for 10 min at 37°C. Both eluent and input samples (50 μg total cell lysate) were analyzed by immunoblotting using anti-SUR1. These experiments were repeated with slight variations in the transfection protocol. For AKAP220 KD experiments 30 nM of AKAP220 or scramble siRNA was used for transfection. The transfections for AKAP150 KD rescue experiments with WT AKAP79-GFP, AKAP79ΔPKA, or AKAP79ΔPIX were performed using 20 μg of AKAP150 shRNAi plasmid DNA and 5 μg of WT AKAP79-GFP, AKAP79ΔPKA, or AKAP79ΔPIX plasmid DNA. In addition to immunoblotting for anti-SUR1, input samples were analyzed using anti-AKAP150, anti-AKAP220, and anti-GFP to confirm transfections were successful.

### Statistical Analysis

All data were analyzed with the program GraphPad Prism™. Results were expressed as mean ± standard error of the mean (SEM). One-way analysis of variance (ANOVA) followed by the post hoc Dunnet’s test or Tukey’s test were used for multiple comparisons as detailed in figure legends. When only two groups were compared, unpaired Student’s t-tests was used. The level of statistical significance was set at *p*<0.05.

## Data availability

All data are contained in the manuscript.

## Acknowledgements

We thank Dr. Christopher Newgard for the rat insulinoma INS-1 clone 832/13 cells. We also thank Dr. Manuela Zaccolo for the cAMP CUTie sensor plasmids and Dr. John Scott for the AKAP150 shRNAi plasmids. Finally, we thank the Advanced Light Microscopy Core at Jungers (OHSU, Portland, OR) staff for their expertise in image acquisition and analysis. This work was supported by National Institutes of Health grant R01DK057699 and 3R01DK057699-14S1 (to Show-Ling Shyng). Dr. Mark Dell’Acqua acknowledges the support of NIH grant NS040701.

## Author contributions

VC designed and performed experiments, analyzed data, and wrote the manuscript; ZY performed experiments; MD provided reagents and edited the manuscript; SLS conceived the project, analyzed data and wrote the manuscript. All authors have full access to all the data in the study and take responsibility for the integrity of the data and the accuracy of the data analysis. All authors reviewed the results and approved the final version of the manuscript.

